# MetaEntropy: quantifying protein variability in complex viral mixtures via entropy

**DOI:** 10.64898/2026.11.17.767350

**Authors:** Leandro Roberto Jones, Julieta Marina Manrique

## Abstract

High-density sequence data has become commonplace in the study of complex viral mixtures from wastewater. Established algorithms are available for lineage deconvolution and labelling of clinically-elusive mutations. However, less attention has been paid to the quantification of translated amino acid diversity. We present MetaEntropy, a modular R package for assessing protein sequence variability and identifying sites under potential selection via Shannon entropy. MetaEntropy integrates utility functions for visualizing genome-wide and protein-specific variability, extracting nucleotide and amino acid mutation profiles, and identifying entropy hotspots. Despite being designed for wastewater surveillance, the package can also profile protein diversity from cross-sectional intra-patient data. However, it does not analyze population structure or quasispecies, for which several tools are available. By focusing on positional entropy, MetaEntropy bypasses the sequence constraints, computational overhead, and consensus-based variant loss typical of traditional haplotype reconstruction and multiple sequence alignment strategies. To balance sensitivity against neutral variation, the package evaluates entropy using a standard 20-class canonical amino acid scheme and a 7-class model based on traditional physicochemical properties. Calculated entropy values are normalized to a 0–1 scale, enabling direct comparisons of protein diversity across loci and sample strata. As proofs of concept, we employ MetaEntropy to detect well-known SARS-CoV-2 functional changes from wastewater data, and to explore tissue-specific heterogeneity in systemic infections. MetaEntropy is available from the Comprehensive R Archive Network (CRAN) at https://CRAN.R-project.org/.

## Introduction

Massively parallel sequencing brought about a qualitative leap in population-level microbial study. While traditional sequencing allowed for tens or hundreds of sequences per locus to be inspected, modern technologies can achieve figures in the tens of thousands. This significantly improved wastewater-based virological surveillance, which relies on sequencing the viruses present in samples of such water [1, 2]. The approach has been successfully used to track previously detected and clinically-elusive variants, for which several bioinformatic tools have been developed [3].

Conversely, wastewater remains underutilized as a source for molecular evolutionary analysis. Maintaining clinic-based genomic surveillance programs is cumbersome even for high-income countries. In this context, wastewater represents an attractive complement to classical patient-centric sampling by aggregating circulating viral lineages within a single matrix. MetaEntropy contributes a tool to estimate circulating protein diversity from translated wastewater sequences. As a metric, it employs Shannon entropy (*H*), which has been widely used in the study of biodiversity and is sensitive to functional variability [4, 5].

While the package’s primary design targets wastewater surveillance, it can accommodate other population-level datasets, such as intrahost variants. However, it does not address specific theoretical aspects of intrahost variation, such as quasispecies dynamics, viral fitness, sequence space occupancy, and error thresholds, which are proper to the quasispecies framework [6, 7].

Table 1 shows a comparison of MetaEntropy with representative tools geared toward the analysis of complex viral populations. The listed wastewater-focused tools (`MixviR` and `lineagespot`) specifically aim at SARS-CoV-2 lineage identification and quantification. It is worth noting that other `R` packages not listed in Table 1 aim to reconstruct epidemic dynamics or forecast clinical incidence using wastewater viral loads, such as `EpiSewer`, `ern`, `WES`, and `wwinference` [8–11]. The listed tools designed to study viral quasispecies generally aim to characterize population structure through haplotype reconstruction, diversity indices, and mutation dynamics. In general, they do not focus on protein-level variation, with the notable exception of `DiversiTools`, which computes amino acid frequencies alongside variant calling. Outside of `DiversiTools`—from which residue-level entropies could be calculated manually with downstream scripting—the remaining applications would require advanced custom programming to yield protein-level metrics. Finally, to our knowledge, no existing applications provide biochemically based functional assessments natively.

**Table 1.** Comparison of MetaEntropy with representative population-oriented tools.

| Tool | Type | Biological Framework | Primary aim | Residue-level analysis | Genomic scope<sup>1</sup> | Amino acid classes | Functional assessment | Input | Platform<sup>2</sup> |
|---|---|---|---|---|---|---|---|---|---|
| MetaEntropy | R package | Wastewater | Predicted protein entropy | Yes | CDS | Yes | Biochemical | SNV table, CDS map | Agnostic |
| MixviR | R package | Wastewater | SARS-CoV-2 lineage profiling | No | Whole genome | No | – | VCF file, reference, BED file, lineage signatures | Agnostic |
| lineagespot | R package | Wastewater | SARS-CoV-2 lineage profiling | No | Whole genome | No | – | VCF file(s), reference, lineage signatures | Agnostic |
| QSutils | R package | Quasispecies | Diversity profiling, mutation dynamics | No | Locus-specific | No | Ecological<sup>3</sup> | Imported alignment and haplotype frequencies | SRS |
| longreadvqs | R package | Quasispecies | Haplotype reconstruction, OTU-based diversity profiling | No | Locus-specific | No | – | Imported alignment | LRS |
| QAP | Stand-alone platform | Quasispecies | Haplotype reconstruction, OTU-based diversity profiling | No | Universal | No | – | FASTQ/FASTA reads, reference | Agnostic |
| QuasiSeq | Pipeline | Quasispecies | Full-length quasispecies reconstruction | No | Whole genome | No | – | Long reads, reference | LRS |
| DiversiTools | Pipeline | Quasispecies | Variant calling, diversity profiling | Yes<sup>4</sup> | Universal | No | – | BAM alignment, reference, CDS map | SRS |

<sup>1</sup>Technically, all tools can take any sequence, but some algorithms are optimized for targeted data. <sup>2</sup>Indicates which technology the algorithms are optimized for, even though they may use other platforms. *SRS* short-read sequencing, *LRS* long-read sequencing. <sup>3</sup>Functional diversity. <sup>4</sup>Reports amino acid frequencies.

## Materials and Methods

### Design

#### Rationale

The design of MetaEntropy addresses the inherent trade-offs in processing viral genetic material from wastewater-based surveillance. While this material is suitable for haplotype reconstruction, achieving both adequate depth and accurate phasing requires hybrid short- and long-read sequencing, which introduces substantial sequence constraints and computational bottlenecks [12]. MetaEntropy minimizes these data processing and financial costs by shifting the analytical focus to positional entropy, thereby bypassing haplotype inference entirely. This design choice also eliminates the need for multiple sequence alignment and prevents the consensus-based variant loss typical of standard pipelines. Additionally, by directly ingesting variant frequency tables, the package seamlessly integrates into existing workflows, delegating preliminary steps such as quality control, reference alignment, and variant calling to specialized, established tools. To calculate positional Shannon entropy (*H*) while balancing analytical sensitivity and robustness, MetaEntropy implements two discrete amino acid classification schemes. The high-resolution scheme treats each of the 20 canonical amino acids as an individual class (*k* = 20), prioritizing sensitivity by capturing every residue substitution as a potential variant. Conversely, the reduced 7-class alphabet aggregates amino acids into six distinct groups based on shared physicochemical properties [4], using a seventh class to accomodate nonsense mutations (Table 2). While the 20-class scheme captures subtle variations, the 7-class physicochemical framework establishes a functional impact threshold by requiring radical transitions (e.g., polar to charged) to register an entropy change, thereby enhancing metric robustness against neutral variations [13, 14].

**Table 2.** Reduced 7-Class Physicochemical Alphabet for Entropy Calculation.

| Class | Amino Acids Included | Physicochemical Properties |
|---|---|---|
| Aliphatic | A, V, L, I, M, C | Non-polar, hydrophobic side chains |
| Aromatic | F, W, Y, H | Bulky, ring-containing structures |
| Polar | S, T, N, Q | Uncharged, hydrophilic side chains |
| Positively Charged | K, R | Basic residues |
| Negatively Charged | D, E | Acidic residues |
| Special | G, P | Conformationally unique residues |
| Nonsense | Stop Codons | Premature translation termination |

#### Input data

MetaEntropy requires data to be provided in two objects with the single nucleotide variants (SNV) identified in the metagenome under study, and the CDS topology of the reference genome used in variant calling (Table 1). The SNV data must be provided in a `data.frame` that includes the frequencies of the reference and alternative bases and amino acids, along with their respective positions relative to the reference genome (Fig. 1). This information is to be distributed in 8 columns labeled, by default, as: `position` (mutated positions), `ref` and `alt` (reference and alternative bases, respectively), `protein` (affected protein names), `aa_position` (mutated protein positions), `ref_aa` and `alt_aa` (reference and alternative amino acids, respectively), and `alt_aa_freq` (alternative amino acid frequencies). The CDS topology (`genome` parameter) must be provided as a `list` with two elements: a `numeric` value (`Length`) giving the length of the reference genome, and a `data.frame`, named `CDS`, informing on gene coordinates and coded protein names. The `CDS` element must have three columns named `start`, `end` and `protein`, containing numeric values corresponding to the first (`start`) and last (`end`) CDS positions in the genome, and the name of each encoded protein (`protein`). An example genome, named `mn908947.3`, is provided with the package. This type of data can be obtained from General Feature Format files using the `rtracklayer` package, or manually, from GenBank or other annotations.

**Fig 1.**
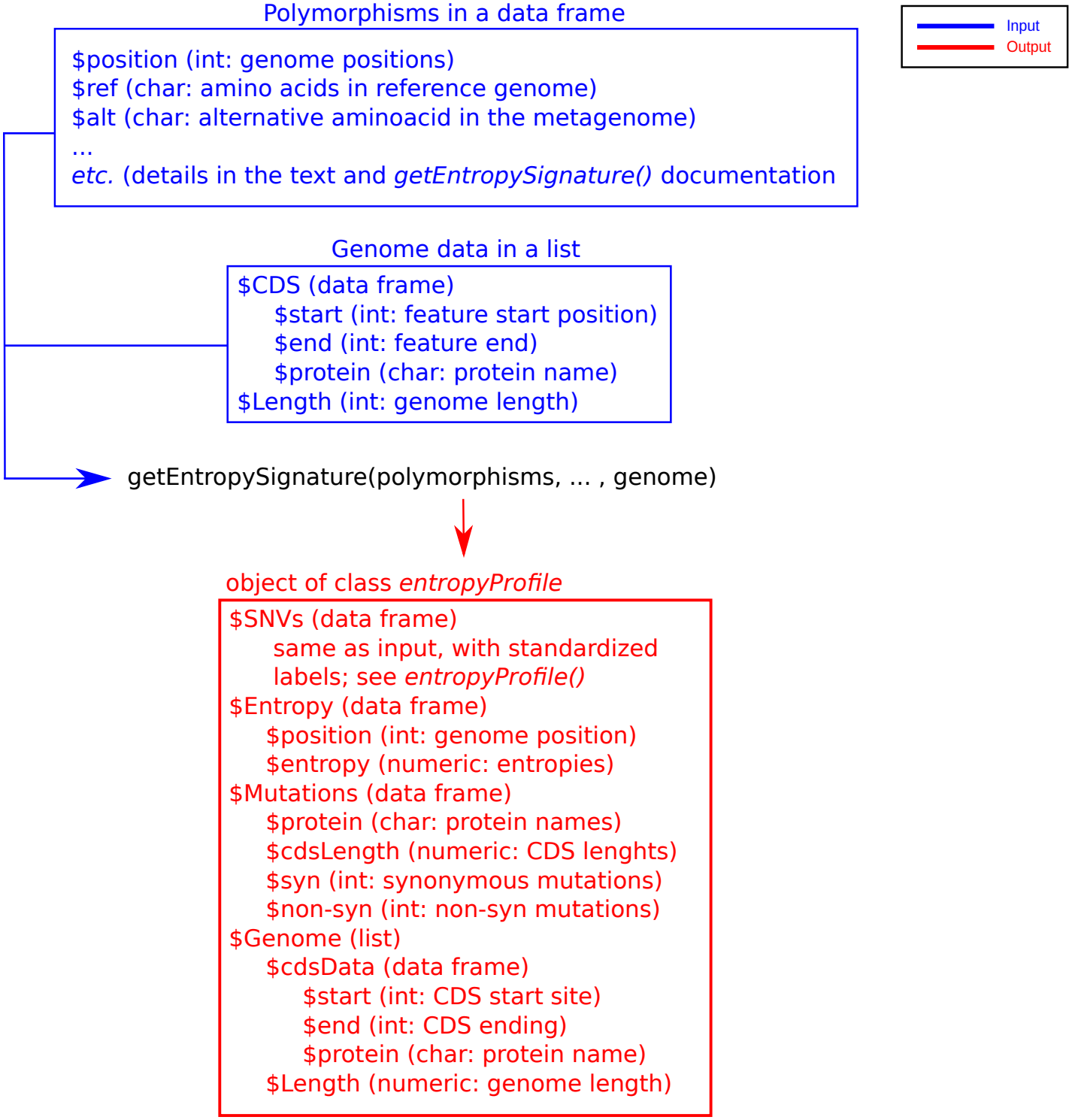
MetaEntropy input and output specifications. Schematic detailing the inputs passed to `getEntropySignature()` and the structural layout of the generated `entropyProfile` object.

Some mutations may be part of a same codon, which can be indicated in a separate column, named `linkage` by default, which provides downstream linked positions relative to mutated positions, or closest upstream positions if there are no downstream positions that are part of the same codon. For example, in the `wWater` dataset distributed with the package, mutations T22673C and C22674T are linked to each other and affect codon 371 of the spike (S) gene:

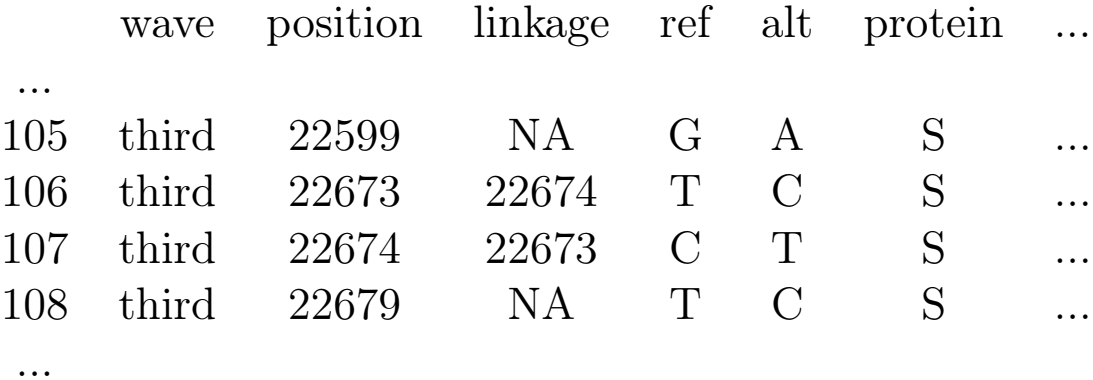

The package does not necessarily need this information since, internally, it focuses on the corresponding amino acid residue. Therefore, multiple substitutions in a same codon can also be encoded using a 3-base notation, conventionally maping the mutation to the second codon base.

In the example above:

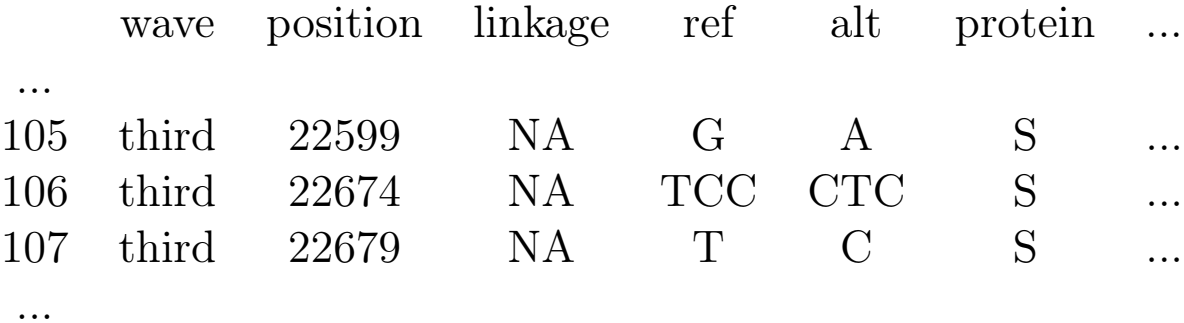

#### Output

Using the `getEntropySignature()` function, the objects described above are processed to orchestrate the *H* computation for an entropy profile (Fig. 1). Entropy values are calculated for each polymorphic codon and mapped to their corresponding protein residue and genomic coordinates. To provide a common coordinate system for variants and lineages with varying genome and CDS lengths, all values are mapped relative to the reference genome. However, the reference itself does not influence the *H* value, as the calculation depends exclusively on variants detected within the virome. The function returns an `entropyProfile` object, which contains the entropy profile, the original input data, and counts of synonymous and non-synonymous mutations per CDS. These latter elements enable visualization and summary statistics, and allow the calculation of normalized mutational prevalences and their comparison with corresponding entropy values, as detailed below.

### Implementation

#### Workflow management

To ensure a modular architecture, MetaEntropy relies on several non-exported internal functions. While these routines are not directly accessible to the user, they are fully documented to facilitate code maintenance and future extensibility.

The input polymorphisms are processed one at a time by `getEntropySignature()`. This function calls `entropyProfile()`, a constructor for the class `entropyProfile`, to create an empty output object. Polymorphisms linked within the same codon exert a combined effect on entropy at the corresponding protein residue. To account for this, `entropyProfile()` allocates a single slot—mapped to the first mutated position of the codon—to store the *H* value shared by these linked positions. Similarly, it maps multiple mutations at the same position to a single entry within the profile. The function also generates the metadata associated with the resulting object, which are required for the user-facing plotting, data extraction, and summary capabilities (detailed below). Input polymorphisms corresponding to synonymous mutations are assigned an *H* value of 0 in the corresponding element of the `entropyProfile` object. Conversely, non-synonymous mutations are processed by `createPositionSummary()`, piped into `fillPosition()`, and finally passed to `getPosEntropy()` to calculate the position-specific entropy (*H*), which is then stored within the `entropyProfile` object. This workflow is schematized in Figure 2.

**Fig 2.**
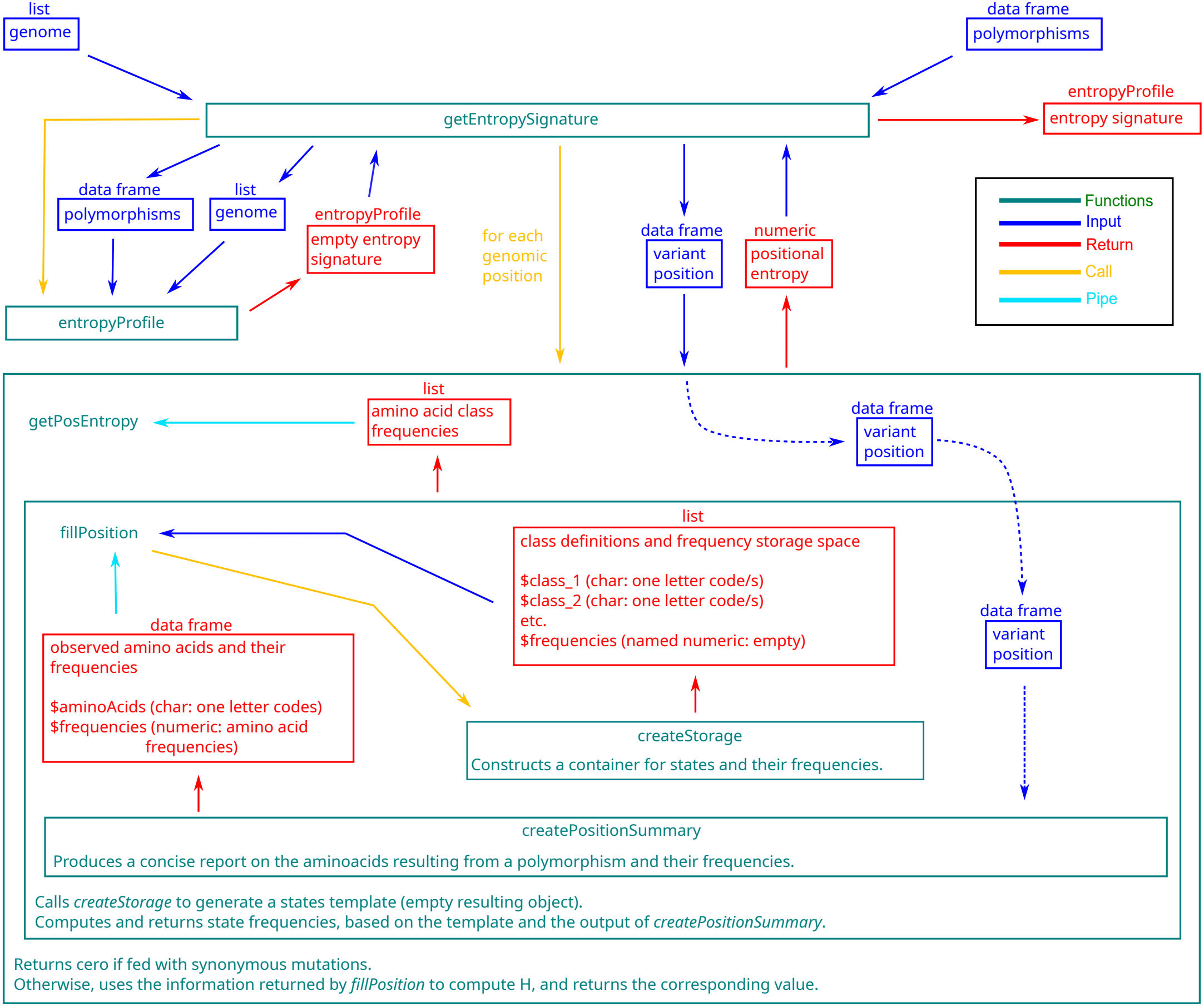
MetaEntropy internal workflow. Conceptual diagram showing the interactions, data structures, and objects passed between internal functions during entropy profile computation.

#### Auxiliary functions

`createPositionSummary()` Notably, each data frame passed to `createPositionSummary()` may contain multiple rows, depending on how many mutations were detected in the virome at the corresponding genomic position. From a biological perspective, multiple haplotypes at a single position indicate increased uncertainty in the protein residue at the population level, reflecting potential functional plasticity and immune evasion. To streamline downstream calculations while preserving this biological significance, `createPositionSummary()` collapses these multi-haplotype dimensions. It condenses each variant position into two vectors, the observed amino acids (a `character` vector of one-letter codes) and their respective frequencies (a `numeric` vector), which are then returned as the columns of a unified data frame.

`createStorage()` The `createStorage()` function initializes the class model structure. Next, `fillPosition()` populates it with class frequencies. Each class description consists of a vector of one-letter amino acid codes. The frequency storage is a named `numeric` vector, where the `names` attribute corresponds to the respective class identifiers. These vectors are returned as a named `list`, providing a structured container for both the class definitions and their corresponding frequency placeholders. This architectural choice ensures the package remains extensible, as downstream functions are designed to process an arbitrary number of classes provided in the aforementioned `list` format. The specific model initialized by `createStorage()` is determined by the `categories` parameter, which is passed to the function via `fillPosition()`.

`fillPosition()` After the generation of an amino acid class structure via `createStorage()`, this function classifies the amino acid variants resulting from a positional polymorphism and calculates the frequencies of the resulting category counts. Its return value is a `list` just like the one created by `createStorage()`, but with the element corresponding to the frequencies of each class filled according to the output of `createPositionSummary()`.

`getPositionEntropy()` The function `getPositionEntropy()` computes positional entropy from state frequencies derived by `fillPosition()`. Using Shannon’s equation, entropy (*H*) is calculated as

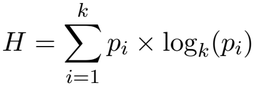

where *k* is the number of amino acid classes and *p*_*i*_ is the frequency of class *i*. To achieve a normalized range of 0 to 1, the function uses the alphabet size (*k*) as the logarithmic base. By the change of base rule, this can be expressed in terms of any arbitrary base (*b*) as

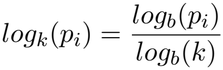

Substituting this into the summation, we obtain

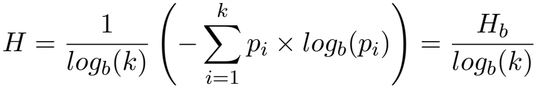

that is the ratio of observed entropy to the maximum possible entropy for that alphabet.

This normalized entropy provides an intuitive scale of functional diversity where *H* = 1 corresponds to a scenario where all functional categories are represented in equal proportions, and *H* = 0 represents the existence of only one functional class. In a biological context, high *H* values are consistent with disruptive or directional selection, while low *H* values reflect stabilization scenarios, such as purifying selection. This standardized formula also allows for the direct comparison of different alphabets; for example, entropy at the genome level with that at the protein level.

#### User-facing Functions

The principal function in the package is `getEntropySignature()` which, as detailed above, is in charge of creating the entropy profile *per se*. This function has nine parameters called `position`, `linkage`, `ref`, `alt`, `protein`, `aa_position`, `ref_aa`, `alt_aa`, and `alt_aa_freq`, used to indicate in which columns of the input are the variant genome position, their linkage relationships, the reference and alternative bases, the mutated protein names, the mutated amino acid position, the reference and alternative amino acids, and the alternative amino acid frequencies in the metagenome, respectively. The default values for these parameters are their own names, but they can be redefined if the raw data labels were to be kept. For example, protein labels are integrated by the `iVar` program into a column that also contains other genomic features, which is named GFF_FEATURE. If this tag was preserved in the polymorphism `data.frame`, the argument `protein=“GFF_FEATURE”` must be passed to `getEntropySignature()`.

The `categories` parameter determines the state scheme to be used in the calculation of *H*. The `sensitive` option sets the classic scheme of one class per amino acid, whereas the `robust` option triggers the functional grouping scheme described in Design section above.

Profiles generated by `getEntropySignature()` can be visualized as per-protein entropy accounts and as genome-wide entropy distributions. For per-protein accounts, the `plot()` method for S3 `entropyProfile` objects can generate strip-, boxplot-, and swarm-style plots (parameter `chartType` set to `stripchart`, `bp`, or `swarm`, respectively). Genome-wide distribution plots (`chartType=“entroScan”`) are scatter plots of H values against genomic position, with points color-coded according to the affected proteins. The default `summary()` function for the `entropyProfile` class produces a table containing per-protein H-value statistics (min., Q1, median, mean, Q3, and max.), along with synonymous and non-synonymous mutational incidences normalized by CDS length. The function `as.data.frame.entropyProfile()` is used to coerce objects of class `entropyProfile` into data frames. It returns a minimal representation of the profile, including the variant genome position (column `position`), the name of the affected protein (`protein`) and the corresponding *H* value (`entropy`).

The package also includes `showMutations()`, a utility function designed to extract and display specific subsets of the SNV table in a more interpretable format. This is useful for identifying specific mutations responsible for high *H* values observed in localized amino acid stretches. The function outputs a formatted table containing three columns: nucleotide-level mutations, their corresponding protein phenotypes, and the alternative amino acid frequencies within the virome. Nucleotide-level mutations are represented by combining the reference base with its genome position and the alternative base (e.g., T22882G). Protein mutations are described by the protein name followed by a colon and a notation consisting of the single-letter reference amino acid code, the amino acid position, and the alternative amino acid code. For example, S:N440K denotes a substitution of asparagine (N) by lysine (K) at position 440 of the spike (S) protein.

To perform graphical and formal analyses of *H* values in specific protein segments (e.g., amino acid stretches with high mutational density), the user is provided with a wrapper for the `boxplot()` and `kruskal.test()` base functions. The function, `assessHotSpot()`, generates various plot types comparing *H* values within the evaluated region against the background distribution of all other protein residues. Additionally, it reports the results of a Kruskal-Wallis test to evaluate the statistical significance of the observed differences. The available visualization styles include boxplots (`chartType=“boxplot”`), strip plots (`stripchart`), and swarm plots (`swarm`).

## Results

In this section we illustrate the use of MetaEntropy by applying it to SNV datasets representative of viral mixtures from wastewater [15] and from systemically infected individuals [16]. Both datasets were derived from purified viral RNA, which was reverse-transcribed to cDNA, enriched via tiled PCR, and subjected to paired-end short-read sequencing. The reads thus generated were aligned to the genome of the Wuhan-Hu-1 isolate of SARS-CoV-2, and the obtained alignments were used for variant calling via well-established programs. The corresponding workflows are briefly described on the `wWater` and `intraHostVariants` help pages. This design, widely used in population studies of viruses, produces exactly the type of data that MetaEntropy is geared towards.

### Wastewater data

The `wWater` data originate from epidemiological weeks 41-2020 and 1-2022 in Argentina. Week 41-2020 was dominated by ancestral lineages, whereas week 01-2022 captured the transition phase as Omicron displaced pre-existing variants to achieve global dominance. We will use MetaEntropy to detect signals of selection in the receptor-binding domain (RBD)—well known for its role in adaptation to humans—during the Omicron era. After applying a minor variant threshold, the code bellow computes entropy signatures using `getEntropySignature()`:

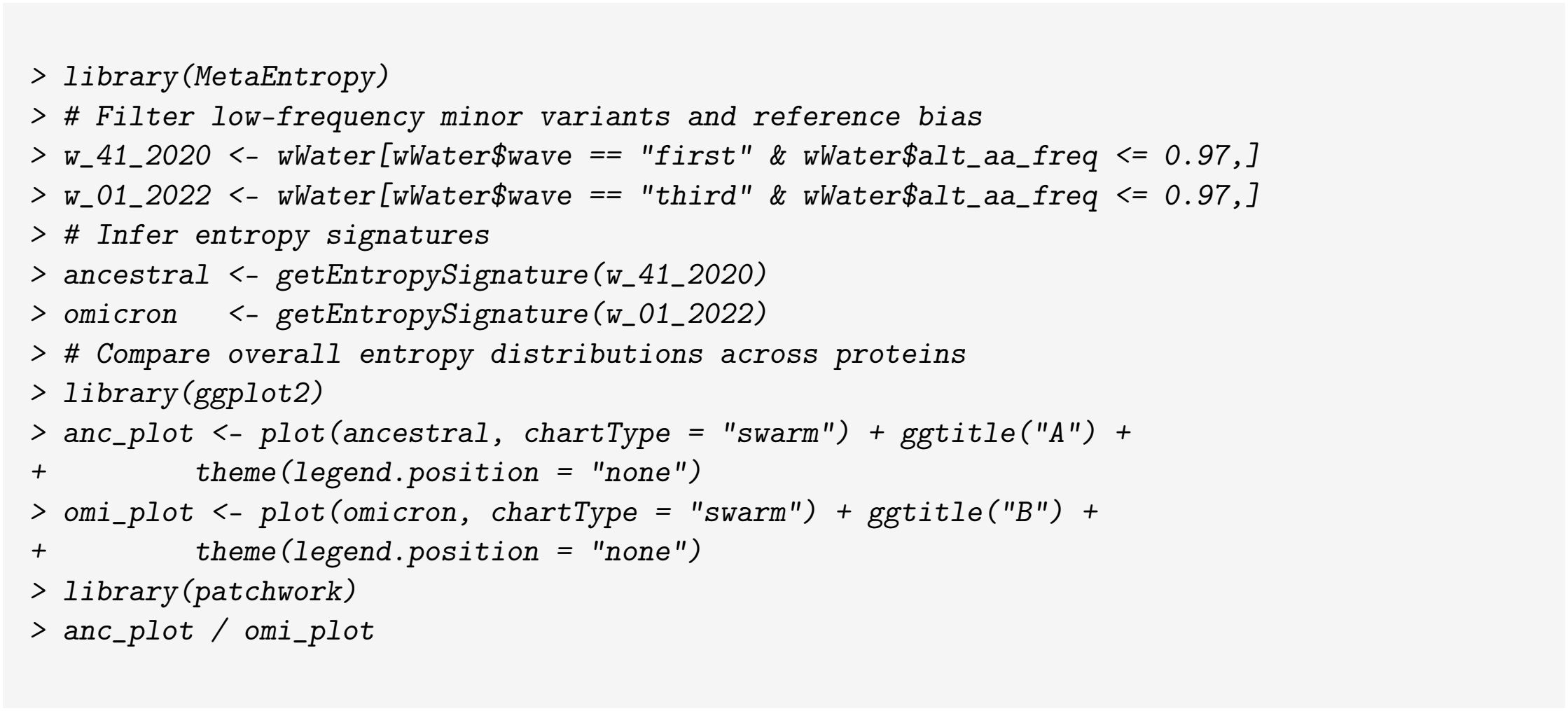

The resulting entropy profiles reveal high entropy concentrated within the Spike (S) protein during the Omicron wave relative to ancestral lineages (Fig. 3). The corresponding amino acid changes can be displayed by using `showMutations()`:

**Fig 3.**
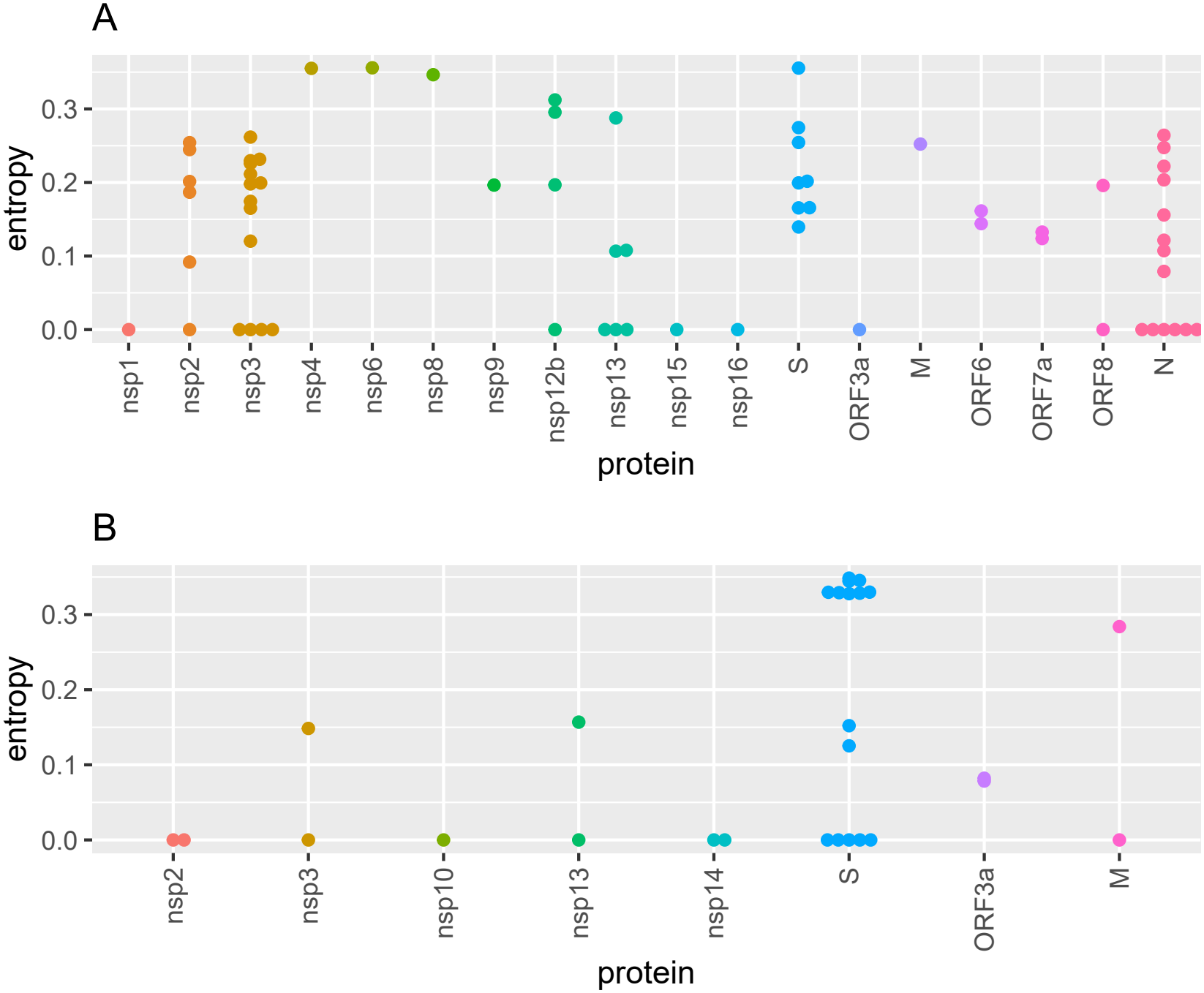
Comparison of entropy signatures between ancestral (A) and Omicron (B) SARS-CoV-2 lineages. Top and bottom panels display entropy distributions across mature viral peptides inferred from non-synonymous SNV frequencies.

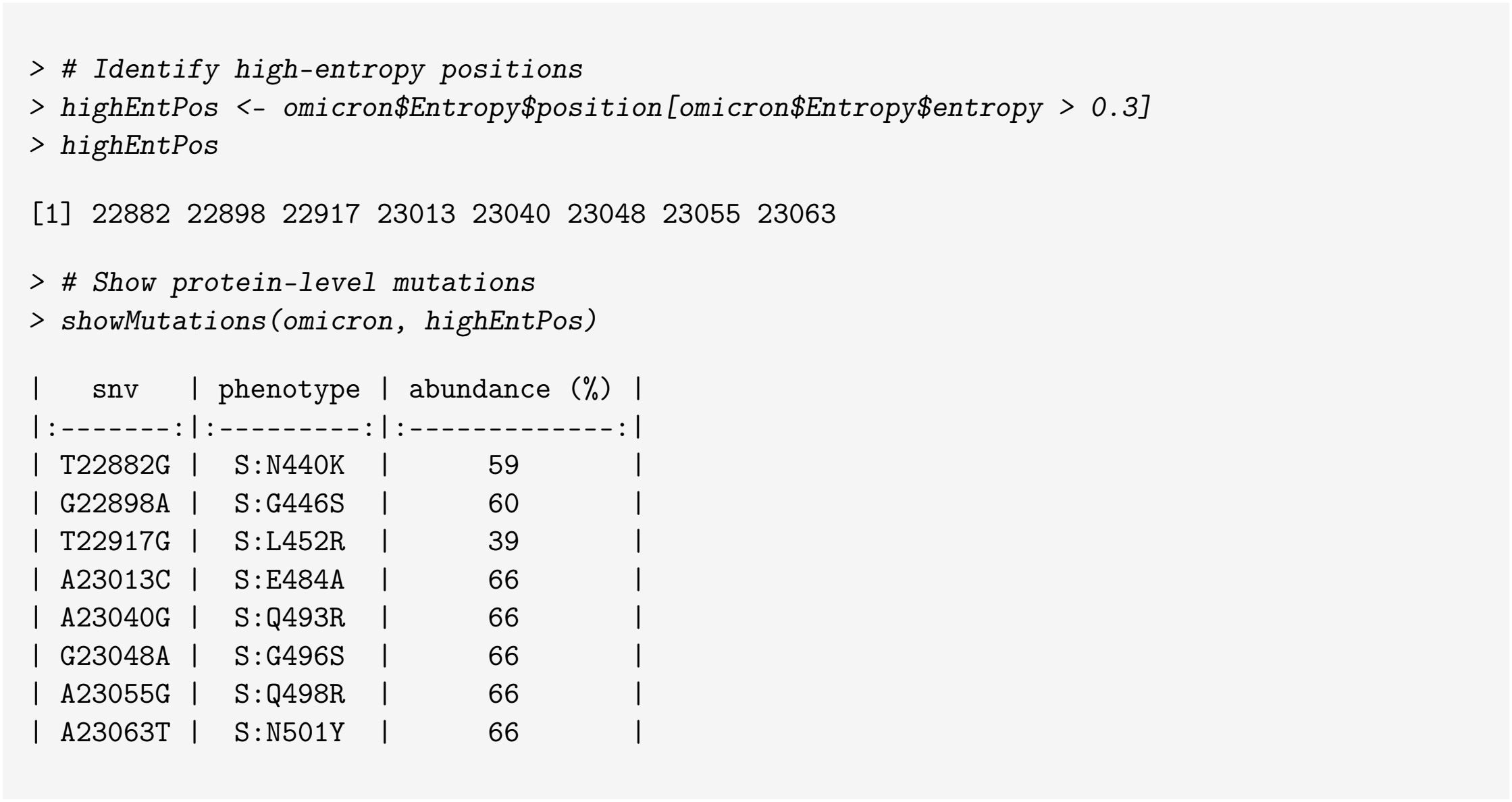

This pinpoints key mutations S:N440K, S:G446S, S:L452R, S:E484A, S:Q493R, S:G496S, S:Q498R, and S:N501Y, located within the RBD (genomic positions 22517–23186). A formal assessment with `assessHotSpot()` confirms that the RBD acted as a significant entropy hotspot during the emergence of Omicron, whereas no significant enrichment was observed among ancestral lineages:

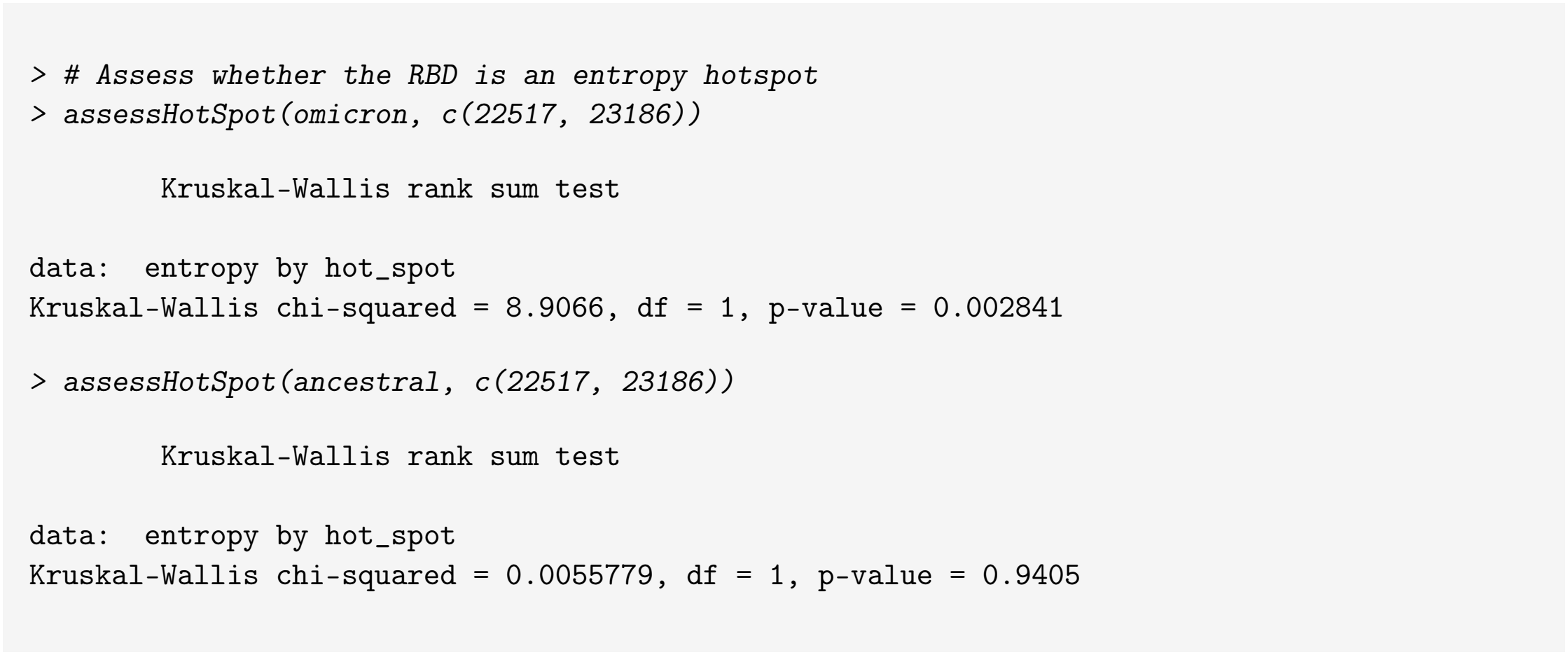

Finally, summarizing entropy signatures using the default `summary()` method for S3 objects of class `entropyProfile` integrates both RNA-level mutational density and physicochemical variation:

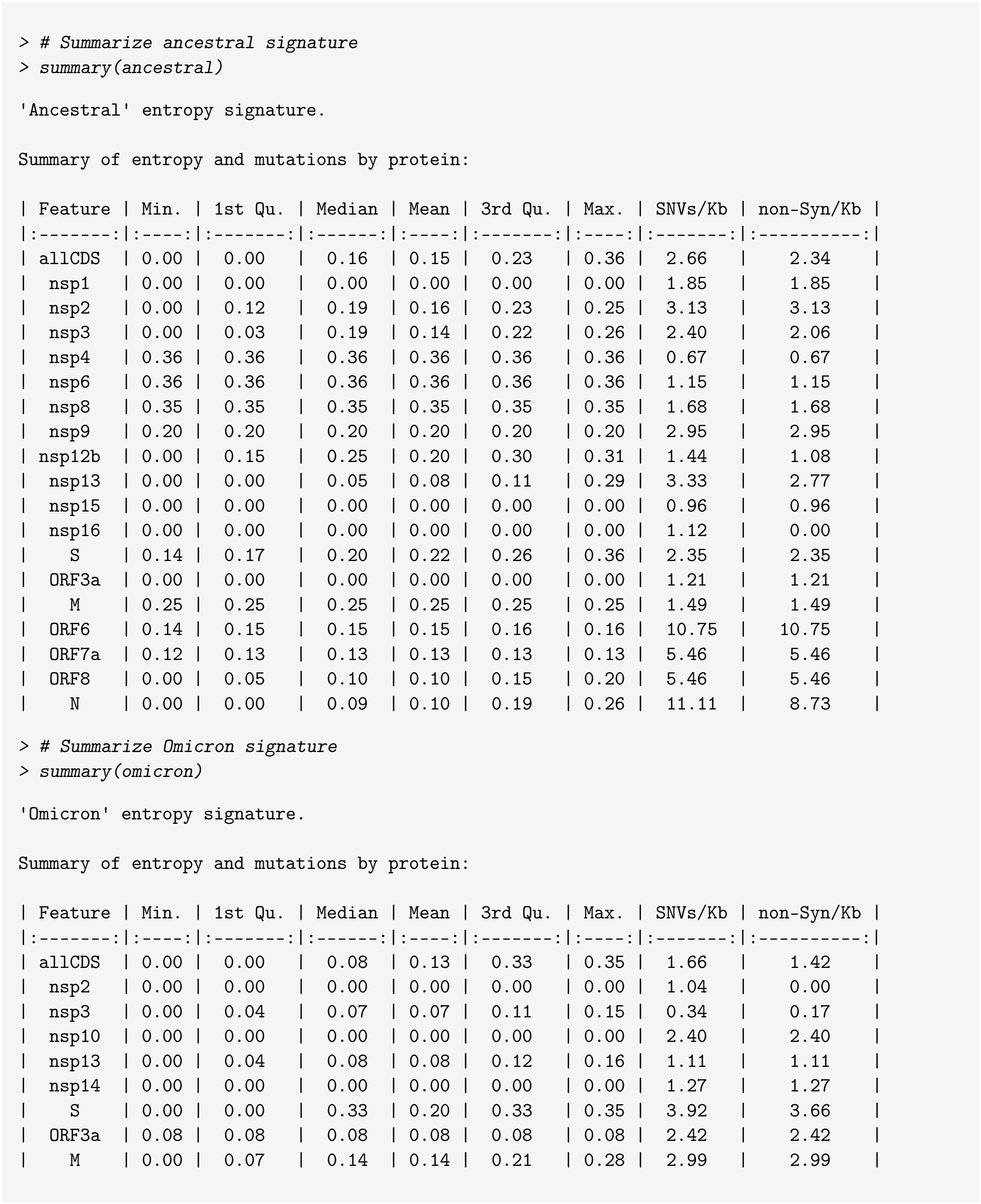

While proteins like N and ORF6 display high raw mutation rates, S exhibits persistently high entropy metrics, particularly during the period coinciding with the global displacement of previous SARS-CoV-2 variants by the Omicron lineage.

### Intra-host variants data

The `intraHostVariants` dataset contains 213 SNVs inferred from four autopsies across five anatomical compartments: lung, intestine, heart, kidney, and liver. We will use MetaEntropy to evaluate whether the proteome at the primary site of infection is representative of the whole within-host diversity.

Because annotations can vary between bioinformatic pipelines, MetaEntropy facilitates custom reference genome modifications. The reference sequence used for generating intraHostVariants is the same as that of `mn908947.3` . However, the nsp12 mature peptide is represented as a continuous block in `intraHostVariants`, whereas `mn908947.3` splits it into its ORF1a- and ORF1b-derived segments. Below we modify `mn908947.3` to create a new `data.frame` to feed the `genome` parameter of `getEntropySignature()`, with nsp12 encoding adjusted from split segments (nsp12a/nsp12b) into a continuous region to match the input table:

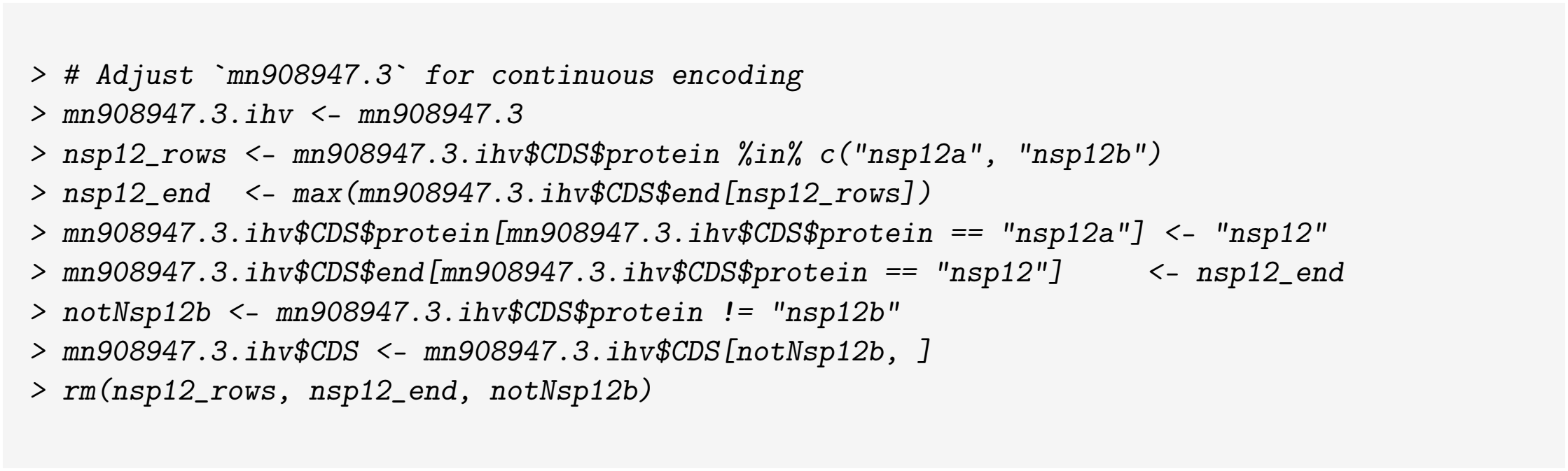

Now we use `getEntropySignature()` to compute entropy profiles for each experimental stratum:

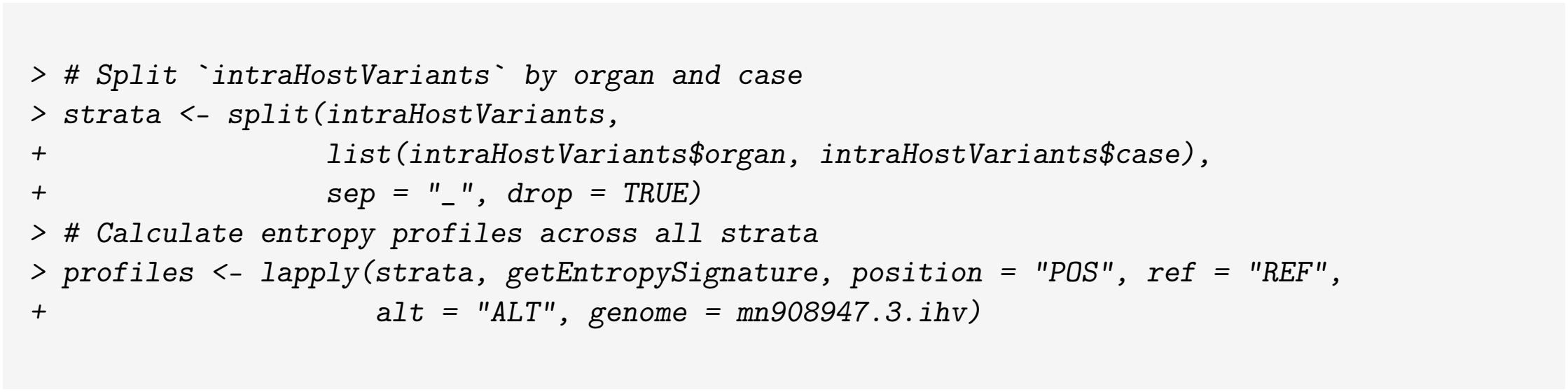

The `heatmap_entropyProfiles()` function allows for graphically comparing multiple profiles. The following code displays variations in mean entropy across tissues and cases:

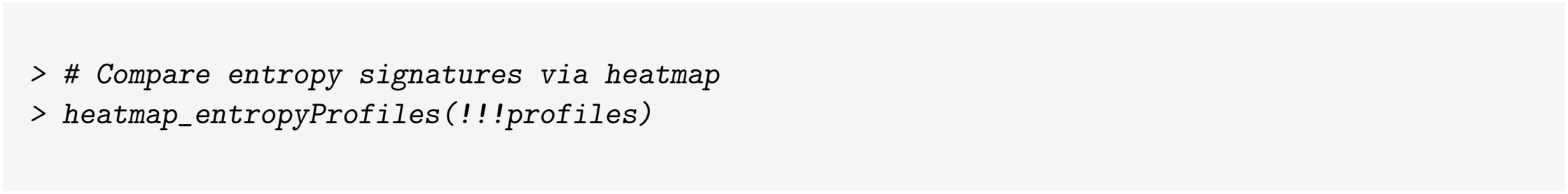

The heatmap indicates that total intra-host viral diversity extends beyond the primary site of infection (Fig. 4). While these results are primarily illustrative, a baseline comparison can be conducted using

**Fig 4.**
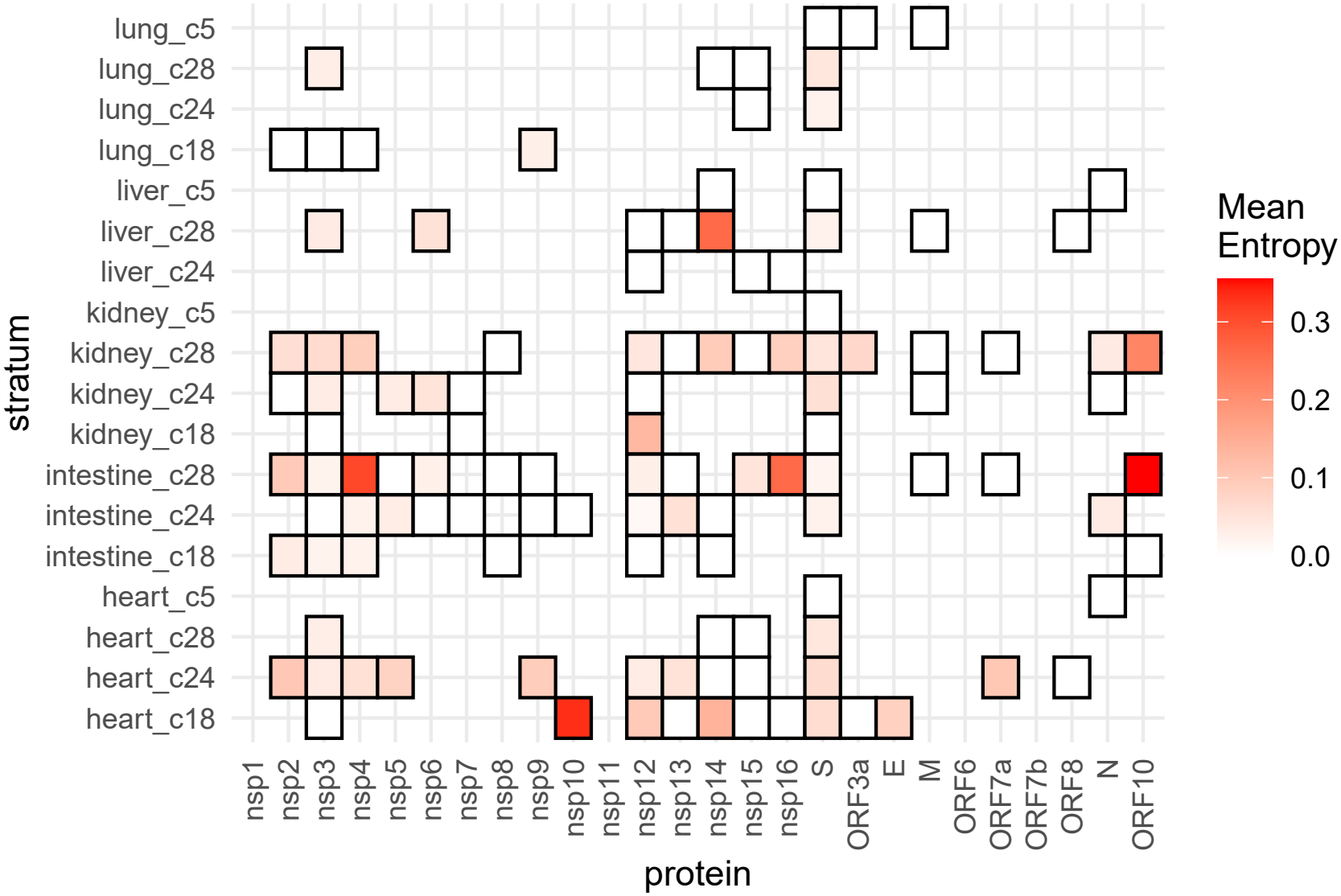
Tissue-specific proteomic variability in systemic SARS-CoV-2 infections. The heatmap displays mean positional entropy across mature viral peptides (x-axis) evaluated across five anatomical compartments (lung, liver, kidney, intestine, and heart) from four autopsies (y-axis).

MetaEntropy’s output. The original study by Manrique *et al*. [16] was designed for an all-versus-all anatomical comparison rather than a respiratory versus systemic assessment. Therefore, evaluating the data under this new binary framework results in an uneven distribution of samples per category (a single lung sample versus a multi-organ pool per individual). To balance detection power, the following code normalizes the data by averaging the entropy indicators across the non-respiratory sites:

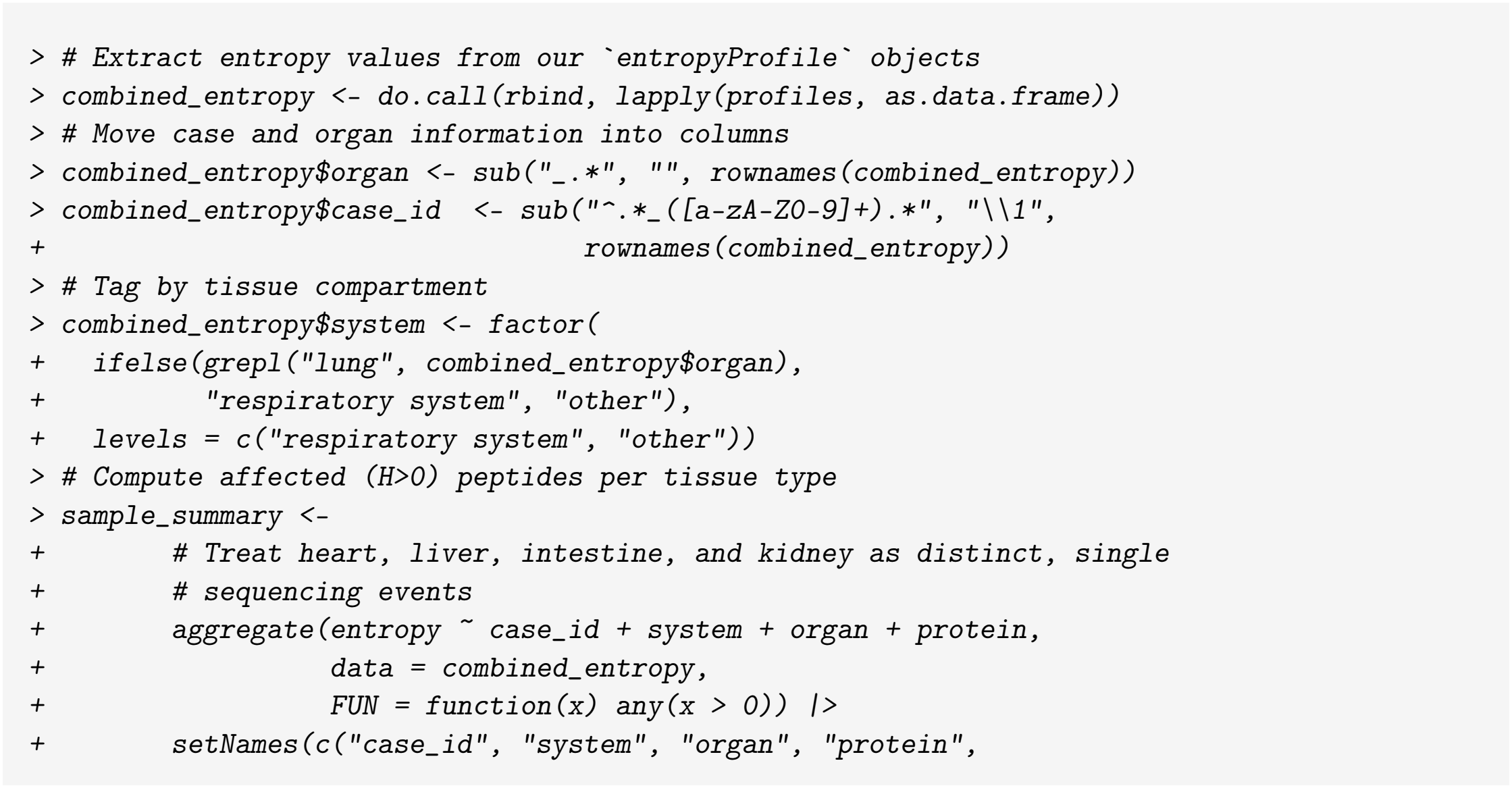
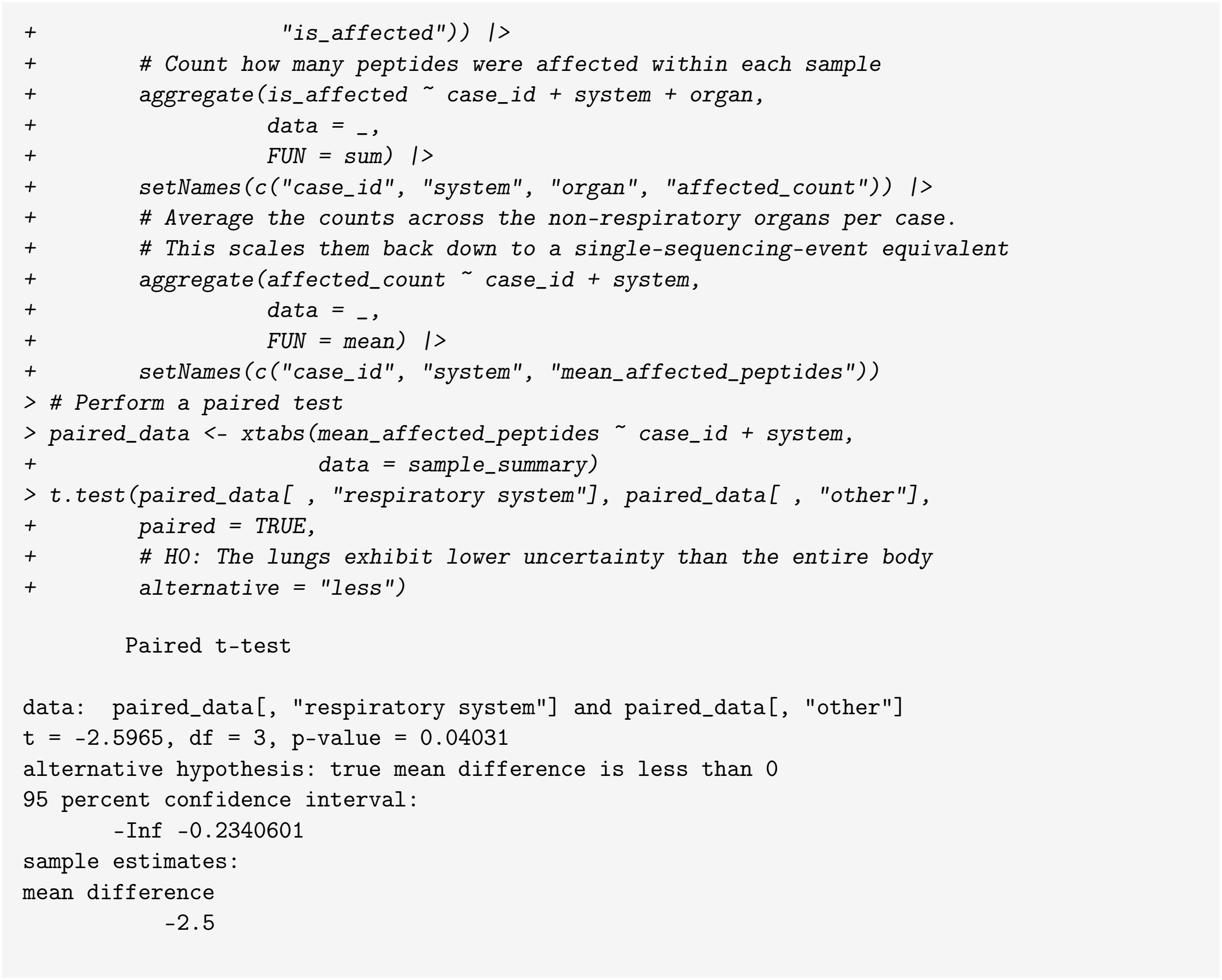

The analysis yields a statistically significant difference in calculated peptide alterations between the respiratory system and the aggregated systemic sites (Fig. 5). Because, as explained, `intraHostVariants` features a sampling aimed at multi-tissue tracking rather than a respiratory-to-systemic contrast, this finding should be treated strictly as a demonstration of the package’s applicability. Nevertheless, the captured patterns are biologically plausible, as discussed below.

**Fig 5.**
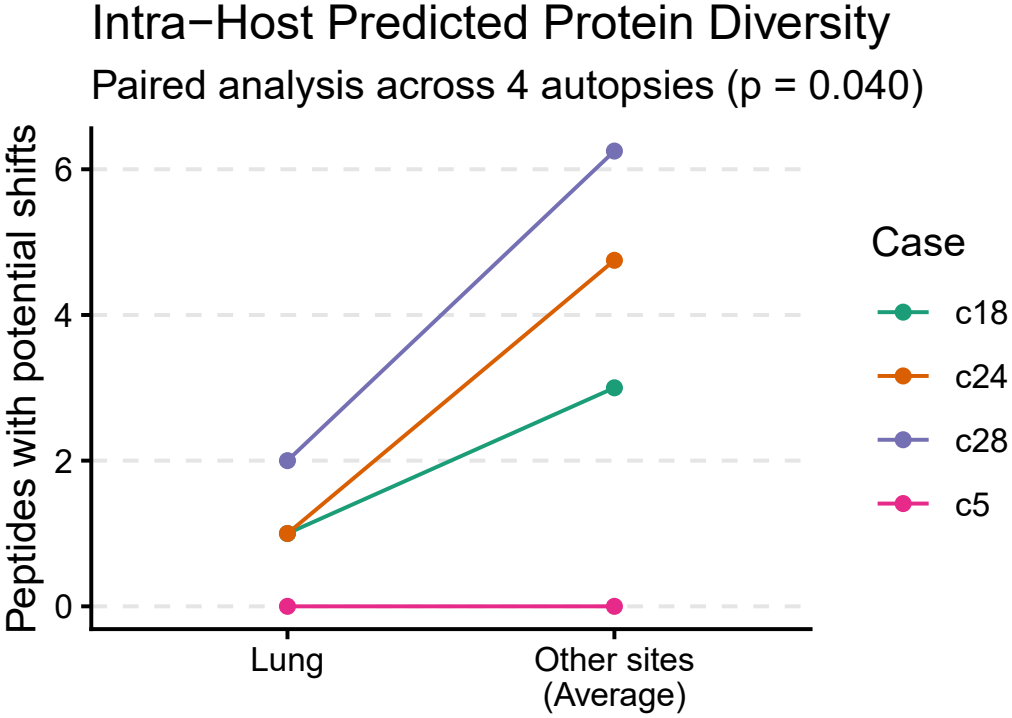
Systemic viral protein diversity in respiratory vs. other anatomical sites. Comparison of unique viral peptides displaying potentially functional changes across patient-matched anatomical sites (*N* = 4 autopsy cases). Each line tracks an individual autopsy, connecting the number of altered peptides found within the primary viral target (respiratory tract) to the average number of altered peptides across all other sampled sites (heart, intestine, liver, kidney). A one-sided paired t-test shows that the observed directional trend is significant (p = 0.040).

## Discussion

The R package MetaEntropy is designed to assess the diversity of protein sequences in complex mixtures of viruses, particularly those present in wastewater. Wastewater, in particular sewage, acts as a sink for the viral diversity in circulation. Therefore, all lineages circulating within a region covered by a collection network have similar probabilities of being detected. As a result, wastewater-based virology presents some interesting characteristics compared to patient-based studies. First, the matrix naturally contains viruses from both clinical and subclinical cases, which is more difficult to achieve in studies based on infected individuals. Additionally, wastewater-based surveillance has the potential to detect new variants before their clinical detection, which in some cases can add important information to early warning systems. On the other hand, traditional datasets often include some sequences that are very closely related to one another. This can result from geographically or socially unbalanced sampling. For example, from biases stemming from the proximity of patients to a research center, or from disparities in access to the healthcare system. Given that wastewater samples represent broad population sectors, its use in epidemiological surveillance, and virological studies in general, help ameliorate this problem. Despite this, and as detailed above, the use of wastewater data in molecular evolutionary analysis has been limited so far. Although high-throughput sequencing yields deep coverage suitable for frequency estimation, fully leveraging these data for protein-level functional insights remains an evolving area. MetaEntropy addresses this by offering a modular workflow that processes codon translations and frequencies derived from standard reference-aligned reads.

The package was designed to provide a measure of variability at the protein level, intended for inferring potential molecular evolutionary processes within the source viral population. In doing so, it addresses a gap in a field where most existing tools focus on lineage profiling and population dynamics. As a metric, MetaEntropy calculates Shannon entropy of amino acid sequences translated from SNVs. This statistic serves as an indicator of functional diversity, as the amino acid side chains in a protein constitutes the primary determinant of its structure and functionality. The package can consider any non-synonymous mutation as potentially disruptive, or alternatively, limit them to radical alterations of side chains—such as the substitution of a small amino acid for a large one, or an aliphatic amino acid for a charged one. This, which aims for greater control over neutral variation, is achieved by implementing a traditional scheme of amino acid classes [4]. It should be noted that a number of other reduced amino acid alphabets have been devised with the aim of capturing functional and structural properties [13]. Some representative schemes are outlined in Table 3. Different reduced alphabets can capture diverse amino acid properties. Furthermore, overlapping schemes can help enhance granularity for the calculation of *H*. As an example of this, let us consider two positions (a) and (b) that exhibit states A and M, and P and Y, respectively, at equivalent frequencies. Under the standard 20-class canonical amino acid scheme *H*_*a*_ = *H*_*b*_. To increase granularity, we could, for instance, assign (a) and (b) to the smallest possible Taylor’s *minimal sets [19],* and the size of that sets could be used to scale *H*. In that way, position (a) would be attributed to Taylor’s *very hydrophobic* minimal set, and position (b) to the *hydrophobic*.*or*.*P* set. In consequence, *H*_*a*_ would receive a scaling factor of 8, and *H*_*b*_ a larger scaling factor of 14, consistent with the greater physicochemical variability observed at position (b) relative to (a). *H* could also be scaled based on the background distributions of amino acid classes. For this, general schemes, such as the BLOSUM62 distribution, could be employed or specific distributions could be inferred for particular viruses—especially for extensively sequenced groups, such as coronaviruses. These are all avenues for enhancing the MetaEntropy’s cappabilities, which will be explored in future versions. Likewise, plans are in place to allow for user-provided reduced aphabets.

**Table 3.** Representative reduced amino acid alphabets.

| Reference | [17]<sup>1</sup> | [18]<sup>2</sup> | [19]<sup>3</sup> | [20]<sup>4</sup> | [21]<sup>5</sup> | [22]<sup>6</sup> | [23]<sup>7</sup> |
|---|---|---|---|---|---|---|---|
| Alanine (Ala, A) | OHyPhi | HyPho | HyPho, Small, Tiny | Ali | Co | HyPhi | STA |
| Arginine (Arg, R) | Ba | VP | P, Ch, Pos, L | Ba | Int | Ba | KREDQN |
| Asparagine (Asn, N) | AcAm | VP | P, Small | Am | Int | HyPhi | KREDQN |
| Aspartate (Asp, D) | AcAm | VP | P, Ch, Ng, Small | Ac | Su | HyPhi | KREDQN |
| Cysteine (Cys, C) | Cys | HyPho | HyPho, P, Small, Tiny | Cys | Co | Slf | C |
| Glutamine (Gln, Q) | AcAm | VP | P, L | Am | Su | HyPhi | KREDQN |
| Glutamate (Glu, E) | AcAm | VP | P, Ch, Ng, L | Ac | Su | HyPhi | KREDQN |
| Glycine (Gly, G) | OHyPhi | MP | Small, Tiny | Ali | Su | HyPhi | G |
| Histidine (His, H) | Ba | MP | HyPho, P, Ch, Pos, Aro, L | Ba | Int | Ba | H |
| Isoleucine (Ile, I) | HyPho | HyPho | HyPho, Ali, L | Ali | Co | Ali | ILV |
| Leucine (Leu, L) | HyPho | HyPho | HyPho, Ali, L | Ali | Co | Ali | ILV |
| Lysine (Lys, K) | Ba | VP | HyPho, P, Ch, Pos, L | Ba | Su | Ba | KREDQN |
| Methionine (Met, M) | HyPho | HyPho | HyPho, L | Ali | Co | Ali | M |
| Phenylalanine (Phe, F) | Aro | HyPho | HyPho, Aro, L | Aro | Co | Aro | F |
| Proline (Pro, P) | OHyPhi | VP | Small | Pro | Int | HyPhi | P |
| Serine (Ser, S) | OHyPhi | MP | P, Small, Tiny | Hx | Su | HyPhi | STA |
| Threonine (Thr, T) | OHyPhi | MP | HyPho, P, Small | Hx | Int | HyPhi | STA |
| Tryptophan (Trp, W) | Aro | HyPho | HyPho, P, Aro, L | Aro | Co | Aro | W |
| Tyrosine (Tyr, Y) | Aro | MP | HyPho, P, Aro, L | Aro/Hx | Co | Aro | Y |
| Valine (Val, V) | HyPho | HyPho | HyPho, Ali, Small | Ali | Co | Ali | ILV |

(1) Based on deduced evolutionary constraints; *HyPho* hydrophobic; *Aro* aromatic; *Ba* basic; *AcAm* acid, acid-amide; *Cys* cysteine; *OHyPhi* other hydrophilic. (2) *VP* very polar; *MP* moderately polar. (3) *Ch* charged; *P* polar; *Pos* positive; *L* large; *Ng* negative; *Ali* aliphatic. (4) *Am* amidic; *Ac* acidic; *Pro* proline; *Hx* hydroxylated. (5) Based on surface-accessibility distribution; *Co* core; *Int* intermediate; *Su* surface. (6) *HyPhi* hydrophilic; *Slf* sulfhydryl. (7) Optimized from Murphy et al. [24] BLOSUM50 clusters to maximize DIAMOND alignment sensitivity against a BLASTX reference.

Using SNV frequency tables, MetaEntropy modularly integrates into workflows featuring established tools for quality control, reference alignment, and variant calling. For instance, in our application examples, fastp and samtools were used for quality control, bwa for reference alignment, and iVar and LoFreq for variant calling. To reduce computational burden, MetaEntropy focuses on positional variability rather than haplotype diversity. This also significantly reduces sequence constraints and avoids the variant loss typical of consensus-based techniques. Despite these advantages, the strategy has an important limitation. By prioritizing positional variability, the tool is blind to the underlying phylogeny; it provides no information regarding the evolutionary history of the observed character-state distribution. This limits the evolutionary conclusions that can be drawn from raw entropy values. Because an ancestral divergence event cannot be distinguished from multiple independent acquisitions, highly entropic sites may simply reflect neutral genetic variation. Therefore, *H* does not serve as a direct indicator of positive selection in the way traditional metrics like *dN/dS* do. As an example, consider a position similar to S:N501Y from the `wwater` dataset analyzed above, where a radical biochemical shift occurs between a small amino acid with a polar side chain and a large aromatic amino acid. While such a change suggests a selective reason, the explanation for the observed distribution of states in our hypothetical example depends on how they originated over time. Although many mutational histories are possible, we illustrate this problem using two simplified scenarios (Figure 6). In the scenario depicted in panel A, the current distribution of states is perfectly explained by a single ancestral change. Conversely, panel B depicts a scenario requiring a minimum of three independent mutational events to explain the terminal distribution. Furthermore, while the change in panel A occurred in a single direction (*N* → *Y* ), the scenario in panel B involves bidirectional substitutions. Under the panel A phylogeny, the current distribution could stem from a single cladogenetic event driven by positive selection, but it could equally be explained by neutral genetic drift where the mutation simply accompanied divergence. Conversely, the panel B scenario strongly implies a sustained selective pressure acting continuously over time to drive recurrent switches between states. This latter pattern is highly characteristic of selection driven by population-level immunity—a common phenomenon in several viral species [25–27]. In summary, heightened entropy does not intrinsically imply ongoing selective forces; rather, it identifies a signature of selection that requires downstream validation through phylogenetic and experimental approaches. Beyond these intrinsic trait, MetaEntropy remains subject to general bioinformatic constraints that affect sequencing workflows, such as PCR biases, systematic sequencing errors, and reference mapping issues. Left unchecked, these technical artifacts can artificially distort the resulting *H* metrics. However, because MetaEntropy integrates modularly into standard pipelines, these extrinsic issues do not represent fundamental limitations of our tool and can be effectively controlled by applying adequate upstream quality control filters and rigorous variant-calling procedures prior to calculating entropy.

**Fig 6.**
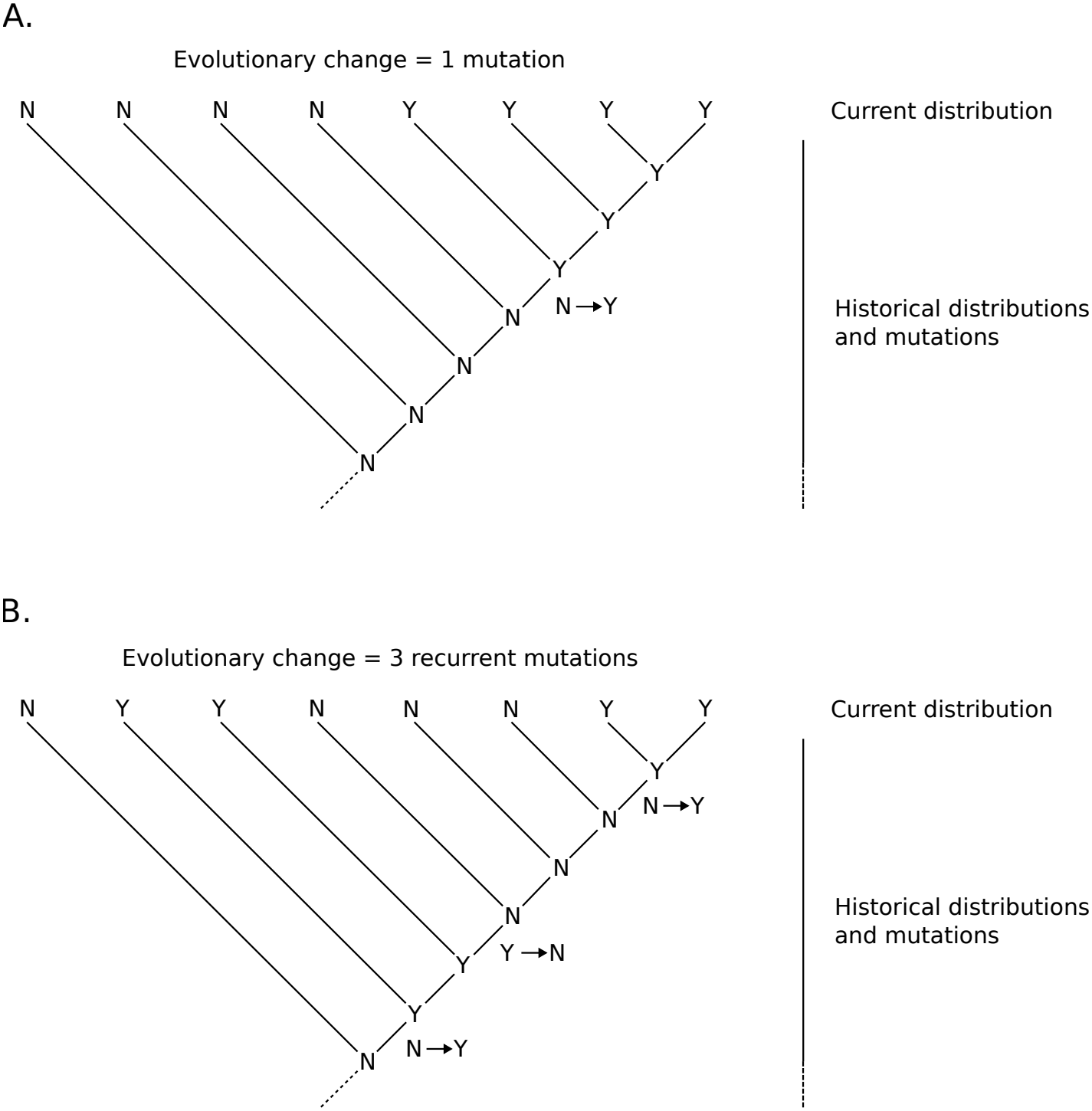
Positional variability versus phylogenetic non-independence in entropy calculations. Diagram demonstrating how identical terminal amino acid distributions can arise from fundamentally different evolutionary histories. States at the terminal nodes represent the current distribution observed within the sequence alignment, while internal nodes are assigned ancestral states. (A) A single mutational event (*N* → *Y* ) accounts for the current distribution. This pattern can be explained either by a selective sweep initiating the clade or by neutral genetic drift accompanying cladogenesis. (B) The identical terminal distribution requires at least three independent, recurrent mutations, strongly implying a continuous selective force sustained over time, such as immune selection. Because MetaEntropy evaluates site-by-site frequencies without tracking tree topology, it calculates an identical value of *H* for both scenarios despite their distinct biological implications.

## Conclusions

MetaEntropy facilitates the extraction of functional insights at the viral protein level directly from SNV tables, providing a modular and streamlined tool that does not require lineage deconvolution, consensus estimation, or multiple sequence alignment.

As shown in the usage examples above, the package captures viral protein signatures across distinct biological scales. When applied to population-level wastewater surveillance, MetaEntropy detected signals of functional change—a phenomenon supported by independent experimental and observational data [28–34]. This highlights the utility of the software for wastewater-based epidemiological surveillance. Analyses of intra-host viral populations using the package revealed potential proteome variations across the body, consistent with the hypothesis that extra-respiratory tissues can act as reservoirs capable of promoting viral evolvability [16, 35–37]. This illustrates how MetaEntropy results can guide investigative directions, such as evaluating whether localized proteomic variation influences immune responses, or determining the extent to which single-site sampling captures whole-host viral protein functionality.

The utility of MetaEntropy lies in its potential as a tool for summarizing and exploring protein sequence diversity. As is the case with other metrics, high values of *H* should not be interpreted as direct evidence of selection. Instead, highly entropic sites should be considered candidates for further investigation using specialized indices, such as dN/dS, and experimental approaches.

## Acknowledgments

This work was partially supported by research grants PIP11220200102657CO from the National Scientific and Technical Research Council (CONICET) and PI1888 from National University of Patagonia San Juan Bosco.

